# Representation-dependent domain adaptation for cross-cohort generalization of mammography diagnostic models

**DOI:** 10.64898/2026.10.01.756041

**Authors:** Ankesh Kumar Pandey, Shandar Ahmad

**Affiliations:** School of Computational and Integrative Sciences, Jawaharlal Nehru University, New Delhi, 110067, India

**Keywords:** Mammography diagnosis, Cohort divergence, batch effects, domain adaptation

## Abstract

**Background:** Artificial intelligence (AI) models for mammography can achieve high diagnostic performance when training and test data originate from similar populations and imaging environments, but performance often deteriorates under cross-cohort domain shift. Differences in acquisition systems, image characteristics, preprocessing, population composition, and disease prevalence can introduce distributional variation that is poorly captured by conventional within-dataset validation. Recent studies have further shown that Foundation Model representations can retain substantial dataset-specific information. However, it remains unclear how image representation influences diagnostic transfer to completely unseen cohorts and whether domain-adversarial learning provides comparable benefits across different representation types.

**Methods:** We systematically evaluated cross-cohort mammography classification using seven independent datasets spanning diverse geographic, institutional, scanner, and imaging environments. Three image representations were compared: handcrafted radiomic features, EfficientNet-B0-derived features, and embeddings from the medical vision-language model MedSigLIP-448. Each was evaluated under three preprocessing conditions: unprocessed images, Otsu-based breast-region extraction, and Otsu extraction combined with Contrast Limited Adaptive Histogram Equalization (CLAHE). Patient-level classification was performed using a mean-pooled multilayer perceptron (MLP) baseline and an attention-based Domain-Adversarial Neural Network (DANN). Generalization was evaluated using a multi-source Leave-One-Domain-Out (LODO) framework in which six datasets were used for model development and the seventh was completely withheld as an unseen test cohort, iteratively across all seven datasets. Performance was primarily assessed by area under the receiver operating characteristic curve (AUC-ROC).

**Results:** Cross-cohort performance depended strongly on image representation and model architecture. MedSigLIP consistently provided the highest performance on unseen cohorts. With MedSigLIP features, mean LODO AUC-ROC increased from 0.598 with MLP to 0.685 with DANN for unprocessed images, from 0.607 to 0.678 after Otsu preprocessing, and from 0.603 to 0.659 after CLAHE+Otsu preprocessing. When DANN–MLP differences were evaluated across held-out datasets and preprocessing conditions, a significant gain was observed for MedSigLIP after correction for multiple comparisons, whereas corresponding gains were not significant for Radiomics or EfficientNet-B0. A mixed-effects analysis similarly showed a positive MedSigLIP × DANN interaction, although this narrowly missed conventional statistical significance. Unprocessed images produced the highest numerical mean performance for MedSigLIP-DANN, but preprocessing itself had no significant overall effect. Notably, linear probing showed strong dataset discriminability in MedSigLIP and EfficientNet representations but little linearly recoverable dataset information in Radiomics, indicating that low dataset separability alone did not predict better diagnostic generalization.

**Conclusion:** Cross-cohort mammography generalization depends strongly on the choice of image representation and its interaction with the classification strategy. MedSigLIP provided the most transferable representation in our multi-source unseen-domain evaluation, and the performance gain associated with DANN was concentrated in MedSigLIP rather than consistently observed across representation types. These findings suggest that the effectiveness of domain-adversarial learning is representation-dependent and that neither conventional preprocessing nor low dataset discriminability alone ensures robust transfer to unseen clinical cohorts.

## 1 Introduction

Mammography remains a central imaging modality for breast cancer screening and diagnosis, and artificial intelligence (AI) based image analysis has increasingly demonstrated the potential to assist radiologists in lesion detection, classification, and risk assessment [1]. Deep learning models can achieve high diagnostic performance when evaluated on data drawn from the same or closely related distributions as their training cohorts [2–4]. However, performance measured under such conditions may provide an overly optimistic estimate of real-world utility. In clinical deployment, mammograms originate from different institutions, scanner manufacturers, acquisition protocols, image-processing pipelines, and patient populations [5–8]. Consequently, models trained in one environment may encounter substantial distributional changes when applied to images acquired in another [8, 9].

Cross-cohort variation can arise at several levels. Differences in acquisition and digitization can alter image intensity distributions, contrast, noise characteristics, resolution, and texture [5–8], while population-level differences can influence breast density, disease prevalence, and the distribution of clinical phenotypes [10–12]. These variations create a domain-shift problem in which the relationship between image features and diagnostic labels may differ across datasets [13, 14]. Importantly, a model can exploit image characteristics that are correlated with the training cohort rather than with the underlying pathology [15]. Such cohort-associated information can provide useful predictive signals during internal validation while reducing performance when the acquisition environment or population changes [16–18].

The problem is particularly relevant to modern pretrained image representations. Foundation Models (FMs) provide high-dimensional visual representations learned from large and heterogeneous image collections and have become increasingly attractive for medical image analysis [19]. Their representations can provide useful transferable features without requiring extensive task-specific training. However, transferability should not be equated with domain invariance. Recent multi-institutional mammography work has demonstrated that FM embeddings can retain substantial information about dataset identity and that task-related and dataset-related information may be entangled within the learned representation [20]. Such findings indicate that high-capacity pretrained representations may simultaneously encode clinically useful visual information and redundant cohort-specific characteristics. Consequently, evaluating performance on aggregated or randomly partitioned datasets may not adequately reveal how a representation behaves when confronted with a genuinely unseen acquisition environment [21–24], a challenge widely documented across other medical imaging modalities [16, 25, 26].

Several approaches have been proposed to improve robustness to domain shift. Image-level preprocessing including intensity normalization, histogram equalization, and breast-region segmentation, can reduce some differences between mammographic datasets [17, 27–29]. However, preprocessing can also modify diagnostically relevant image information or introduce artificial structures, and its effect on external generalization may depend on the underlying representation [8, 30]. At the representation level, “domain-invariant learning” attempts to suppress information associated with the source domain while retaining information relevant to the prediction task. Domain-Adversarial Neural Networks (DANNs), for example, introduce a domain-discrimination objective during model training and use adversarial optimization to discourage the learned representation from encoding dataset-specific information [31]. Although such approaches are intuitively attractive for heterogeneous medical imaging, their effectiveness may depend strongly on the characteristics of the representation to which they are applied [32–34].

Recent work has also highlighted an important methodological issue in evaluating multi-institutional medical AI. Aggregating datasets during model development can conceal substantial differences in performance between individual cohorts, even when overall performance appears satisfactory. This has motivated analyses of dataset-specific errors, representation structure, dataset recoverability, and shortcut-like behavior in Foundation Models [20]. Such studies provide important evidence that dataset identity can remain detectable within learned image representations. However, identifying dataset-specific information and demonstrating improved diagnostic transfer are related but distinct objectives. A representation may contain cohort-associated information without necessarily exhibiting poor diagnostic transfer, while a domain-adaptation method that reduces dataset predictability does not necessarily guarantee improved performance on an unseen clinical cohort.

An important unresolved question is therefore how the effectiveness of domain-adversarial learning depends on the underlying image representation when evaluated under a strict unseen-domain diagnostic task. Handcrafted radiomic descriptors [1, 35, 36], conventional convolutional neural-network features [2–4], and representations derived from large pretrained medical vision models [37–39] capture different aspects of mammographic image structure and may respond differently to changes in acquisition conditions and population characteristics. Similarly, preprocessing strategies that improve apparent image standardization may not necessarily improve the transfer of pathology-relevant information across cohorts. A systematic comparison of these factors within a common experimental framework is therefore needed.

In the present study, we investigate cross-cohort generalization of mammography diagnostic models using seven independent datasets representing heterogeneous geographic, institutional, scanner, and imaging environments. We compare three complementary forms of image representation: handcrafted radiomic features, EfficientNet-B0-derived convolutional features, and MedSigLIP-448 (ViT) embeddings. For each representation, we evaluate three preprocessing regimes: unprocessed images, Otsu-based breast-region extraction, and Otsu extraction combined with CLAHE. We further compare conventional multilayer perceptron (MLP) classification with Domain-Adversarial Neural Network (DANN) training.

To distinguish genuine external generalization from performance arising from shared dataset characteristics, we employ a multi-source Leave-One-Domain-Out (LODO) evaluation framework. In each experiment, six datasets are used for model development while the seventh dataset is completely withheld and treated as an unseen target domain. This procedure is repeated across all seven cohorts, producing a systematic cross-cohort evaluation rather than a single source-to-target experiment. The framework allows us to examine not only whether domain-adversarial learning improves performance, but also whether its benefit is consistent across fundamentally different image representations and preprocessing strategies.

The study addresses two principal questions; First, which image representation provides the greatest resilience to cross-cohort variation in mammographic diagnosis? Second, does domain-adversarial learning improve generalization to previously unseen cohorts consistently across different representations and preprocessing conditions? By jointly evaluating representation, preprocessing, and domain-adaptation strategy under a common multi-source unseen-domain protocol, we aim to distinguish improvements in conventional classification performance from improvements in genuine out-of-distribution robustness.

## 2 Materials and Methods

### 2.1 Data Preparation

In this study the datasets utilized are; CBIS-DDSM [40], MIAS [41], INBreast [42], CMMD [43], RSNA [44], VinDR Mammo [45] and IMDB [46]. The main criteria for dataset selection was, that the cohort prepared should be manageable for us to handle computationally while having sufficient diversity on the level of demography (US, UK, Portugal, China, Australia, Vietnam and India), scanner vendors (Hologic, Siemens, GE etc) and modality (Scanned Films and Full-Field Digital Mammography); to make the test of generalization robust. Except for the IMDB dataset, the rest of six datasets have been studied extensively in the previous mammogram analysis studies [8, 9, 18, 47–52]. Exploratory data analysis of the acquired datasets has been discussed later in the results.

### 2.2 Data Preprocessing

#### 2.2.1 Format Conversion

Preprocessing of the acquired images included conversion of the images from the standard radiological image file format DICOM (.dcm) and PGM (.pgm) to PNG (.png) [22, 53]. This reduced the storage overhead of DICOM’s embedded metadata and avoided the PGM format which is progressively getting obsolete.

### 2.3 Cropping and Filtering Strategies

Mammograms frequently contain prominent non-anatomical artifacts, such as text labels, adhesive tape marks, laterality indicators (e.g., “RCC” or “LMLO”), and scanner-specific processing signatures [54–57]. To eliminate these confounding features and isolate the breast parenchyma, we opted for a multi-stage preprocessing pipeline integrating Gaussian smoothing, Otsu’s adaptive thresholding [58], morphological filtering, and largest connected component (LCC) extraction.

First, to reduce high-frequency acquisition noise and stabilize the subsequent thresholding step, each image was smoothed using a 5 × 5 Gaussian kernel with standard deviation *σ* = 1.1; a well documented step in mammogram preprocessing [59, 60].

Following smoothing, we computed the optimal Otsu global threshold *t*^∗^ on the pixel intensity histogram. Because the breast skin-line boundary typically exhibits a gradual intensity decay that fades into the dark background, standard thresholding can erroneously clip peripheral breast tissue. To preserve this boundary, the operational threshold was offset by 20 gray levels and clamped to the valid range:

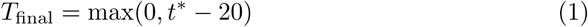

An initial binary mask was constructed from this adjusted threshold, where pixels at or above *T*_final_ were designated as foreground.

To remove persistent artifacts such as labels and lines that survive thresholding, we execute a morphological opening operation using a 5 × 5 flat, square structuring element *B*. The opening of a binary mask *A* by *B* is defined as:

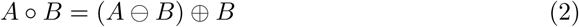

where ⊖ and ⊕ denote the classical morphological erosion and dilation operations, respectively [61]. To eliminate larger extraneous artifacts, this opening was executed for two successive iterations. Because the structuring element is a filled square, two iterations of opening are mathematically equivalent to a single opening using a 9 × 9 structuring element.

Finally, to address any residual disconnected artifacts, we computed the Largest Connected Component (LCC). All isolated foreground regions were treated as independent contours *C_i_*. The optimal segmented breast mask *S*_LCC_ was selected as the contour maximizing the spatial area:

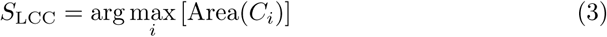

The selected LCC contour was solidly filled to patch small internal voids, and all other contours were discarded, in line with standard mammographic CAD practices [62–64]. The resulting binary mask was multiplied pixel-wise against the original mammogram, preserving the isolated breast parenchyma while mapping all background areas and artifacts to pure black.

### 2.4 Contrast Enhancement

To homogenize intensity distributions and balance localized contrast variations across heterogeneous scanner modalities, via OpenCV we applied Contrast Limited Adaptive Histogram Equalization (CLAHE) [29]. We utilized a tile grid configuration of 8 × 8 and a clip limit parameter of 4.0. These hyperparameters were determined experimentally by optimizing enhancement quality across the aggregated dataset pool. Although modern deep-learning adaptations exist, classical CLAHE remains the robust clinical standard for digital and screen-film mammogram normalization [27, 65, 66].

### 2.5 Image Representation and Featurization Details

We featurized the images using 3 different methods; Quantitative Radiomic features (PyRadiomics package [67]), CNN (EfficientNet-B0 [68]) based latent features and ViT based (MedSigLIP-448 [69]) embeddings. The data cohorts were made compatible for ViT and Quantitative feature extraction by tiling the images by calculating a dynamic sliding window, with the average stride length across all the images under investigation turning out to be 332.13 ± 104.61 pixels. CNNs capture short range dependencies better, so they require uninterrupted spatial maps or whole images to extract the features meaningfully, hence the same were used for CNN based feature extraction [51, 70].

#### 2.5.1 Vision-Transformer based Features

We used “MedSigLIP-448”, a 448 × 448 resolution vision encoder derived directly from the 400M parameter SigLIP (Sigmoid Loss for Language-Image Pre-Training), image-text pair model [71]. We applied it via the ‘Transformers’ library from Hugging Face. Patches with a 90*^th^* percentile intensity below 30, effectively empty background, were filtered out before extraction to prevent unnecessary computation on uninformative tiles. Each valid patch produced 1,152 dimensional embeddings. The training set of MedSigLIP comprises of 32.6 million histopathology patches alongside 635,000 images from radiology, dermatology, ophthalmology, and general medical imaging while lacking Mammograms. This model, to the best of our knowledge, has not been utilized for analyzing mammograms.

#### 2.5.2 Convolutional Neural Network based Features

We selected the EfficientNet-B0 as a global feature extractor. It particularly has shown superiority over other state of the art CNNs in classification task involving chest X-rays [72] and other variants (B2 and B5) have been used in feature extraction of mammograms as well [17], so this provided a strong basis to directly go with this model for extracting CNN based latent features. The architecture uses mobile inverted bottleneck (MBConv) blocks and squeeze-and-excitation optimization and contains 5.3 million trainable parameters. We initialized the network with ImageNet-1K V1 weights, in par with the original architecture and applied ImageNet normalization statistics to ensure consistent feature scaling. The terminal classification head was replaced with an identity mapping. This substitution allowed us to extract 1, 280 dimensional feature vectors from the final global average pooling layer.

#### 2.5.3 Radiomic Features (PyRadiomics)

We extracted a 279 dimensional signature from each patch using the PyRadiomics package. The base set contains 93 descriptors. These included 18 first-order statistics (mean, median, standard deviation, variance etc) along with 5 structural metrics derived from the Neighborhood Gray-Tone Difference Matrix (NGTDM) and four gray-level matrix families, Gray Level Co-occurrence Matrix (GLCM) (24), Gray Level Run Length Matrix (GLRLM) (16), Gray Level Size Zone Matrix (GLSZM) (16) and Gray Level Dependence Matrix (GLDM) (14) – totaling to 93. We calculated this 93 feature set across the original image patch and two wavelet-decomposed sub-bands yielding 279 features per patch. Extraction parameters used a fixed bin width of 25. We applied patient-level Z-score normalization to the extracted features to reduce intra-patient and inter-batch variance.

These three feature extractions, EfficientNetB0, PyRadiomics and MedSigLIP-448 were each run across all three preprocessing cohorts (Unprocessed, Only Otsu, CLAHE + Otsu), producing total 63 (7 datasets × 3 preprocessings × 3 feature types) feature-label files in flat HDF5 format for efficient storage and I/O while training the classifier models.

### 2.6 Classifier Model Details

In the training phase to align with clinical diagnostic workflows, where a diagnosis is reached at the patient level rather than the image level, we adopted a “Multi-Instance Learning (MIL)” setup where each patient id (thus patient) is treated as a bag containing four instances, one for each standard screening view (RCC, RMLO, LCC, LMLO), which further had different patches in our case. For doing the classification we trained a 3 layer simple MLP and a Domain Adversarial Neural Network [31] detailed in each of their own subsections.

#### 2.6.1 Multi Layer Perceptron (MLP)

To establish a baseline of learning, we utilized a fully-connected 3 layer Multi-Layer Perceptron (MLP). To create fixed-length inputs for feeding them in the network, an unweighted mean-pooling operation across all instance-level feature vectors belonging to a single patient was done. The input layer was configured to accommodate the varying dimensionality of these aggregated feature sets: 1,152 for MedSigLIP, 1,280 for EfficientNet, and 279 for PyRadiomics.

The internal architecture consisted of two hidden layers containing 128 and 16 neurons, respectively, each followed by a Rectified Linear Unit (ReLU) activation and a dropout layer (dropout rate = 0.3). The network terminated in a single output neuron generating an raw log odds score (logit). During training, the patient-level logits were optimized using the Binary Cross Entropy loss function. The optimizer used was Adam [73] regularized by combining L1 (1 × 10^−5^) and L2 (1 × 10^−4^) norms with a learning rate of 0.001 and batch size of 32. Class imbalance was managed by incorporating class weight calculations.

5-folds cross-validation for 150 epochs was a standard in training the MLP. Early stopping was utilized with a patience of 15 epochs monitored against the validation set AUC-ROC. During evaluation, logits were transformed into probabilities via a sigmoid activation function, and the optimal diagnostic threshold was determined by maximizing Youden’s J statistic (*J* = Sensitivity + Specificity − 1; *J* ∈ [−1, 1]) [74] on the validation set.

#### 2.6.2 Domain Adversarial Neural Network (DANN)

To enable batch invariant breast cancer detection, a Domain Adversarial Neural Network (DANN) [31] with Multiple Instance Learning (MIL) was used. In this 4-component model, a “Feature Extractor (FE)” network is jointly optimized for the primary classification task (Benign vs Malignant) via the “Task Classifier (TC)” network while being trained to make the learned representation uninformative about the underlying domain via the “Domain Discriminator (DD)” network. A “Gradient Reversal Layer (GRL)” reverses the gradient backpropagating from the DD but leaves the input unaltered by multiplying it with an identity matrix in the forward pass, thereby encouraging the FE to retain **task-relevant information while suppressing domain-specific information**. During our trainings the FE initialized by accepting varying input dimensions depending on the feature type (1,152 for Med-SigLIP, 1,280 for EfficientNet, and 279 for PyRadiomics). It had 2 hidden layers with 512 nodes each, sequentially coupled with Layer Normalization [75]. The TC had a “Gated Attention layer” [76–79] which assigned a normalized attention score to each instance, quantifying its pathological relevance. These scores regulated a weighted sum pooling operation, ensuring that only highly suspicious patches drive the final representation while patches of benign tissue are suppressed to near-zero values. The DD functions exclusively to identify the source dataset of the input features, thus utilized unweighted mean pooling done on incoming feature vectors.

The adversarial dynamics of the architecture came into effect during backpropagation. The TC loss (*L_cls_*) propagated normally to optimize the shared FE for pathology identification. Conversely, the DD loss (*L_dom_*) encountered the GRL, which multiplied the returning gradients by a negative scalar (−*α*). This reversed gradient propagated to penalize the FE for learning batch-specific variations. This mechanism initiated an adversarial mini-max game between the two pathways, forcing the shared FE to simultaneously minimize the *L_cls_* and maximize the *L_dom_*. Consequently, the architecture learned the domain-invariant representations based strictly on features carrying the pathology signal, discarding batch-level artifacts. Figure 1 shows the model architecture with the discussed training mechanism.

**Fig. 1:**
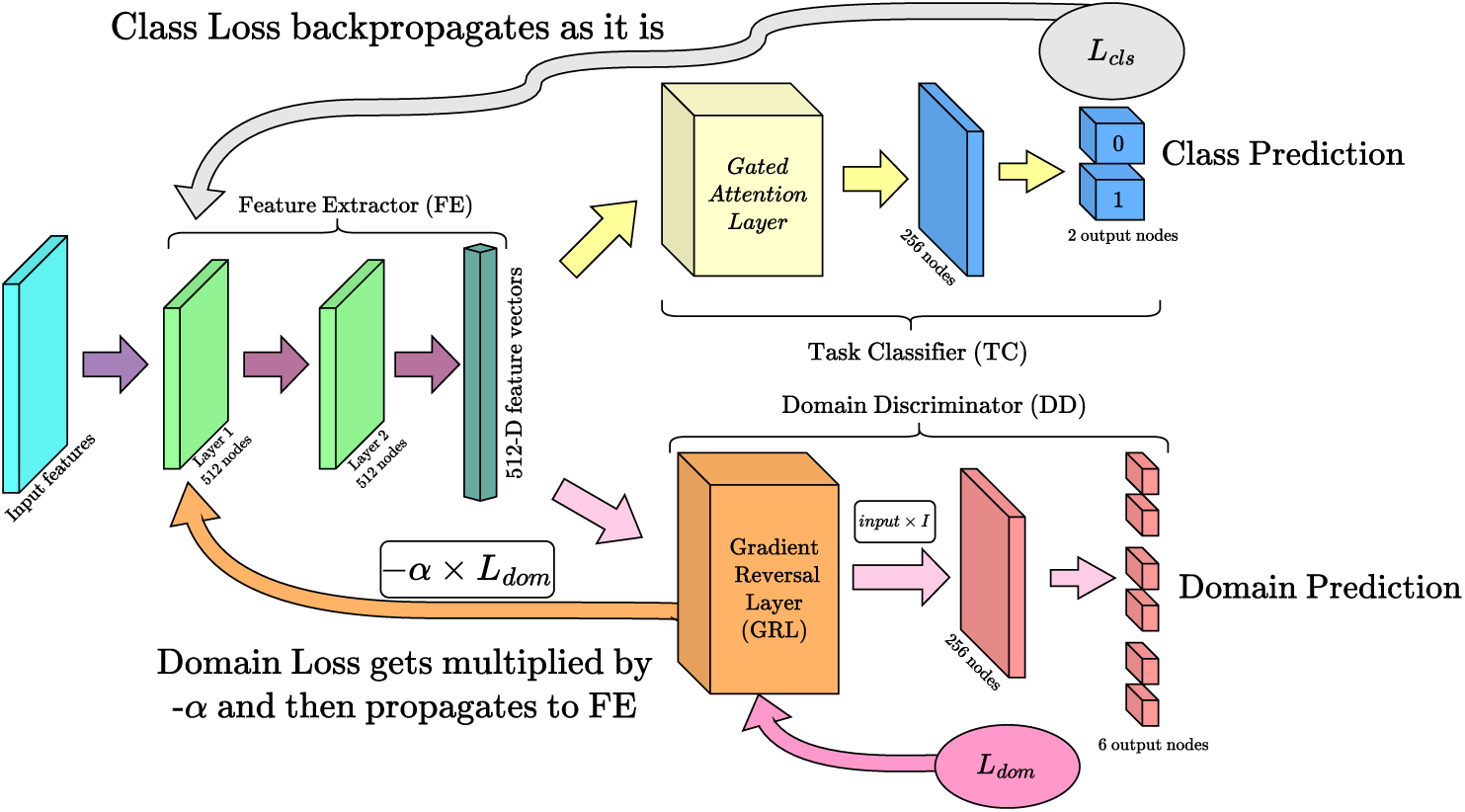
Schematic of the DANN architecture. The diagram follows the conceptual and design layout of the model architecture from the original Ganin et al. paper [31], with components and configurations made specific to the present implementation.

##### DAkk Training Regime

The network was trained using a 5-folds cross-validation don on the training set created after doing patient level stratified shuffle split, running for a maximum of 60 epochs per fold with an early stopping patience of 12 epochs monitored against the validation AUC-ROC. Adam optimizer was used with a learning rate of 1 × 10^−4^, an L2 weight decay of 1 × 10^−4^, and a batch size of one patient bag. In all the three networks (FE, TC and DD) ReLU activation was used and a dropout of 0.5.

The composite loss function summed the cross-entropy classification loss (*L_cls_*) and the cross-entropy domain loss (*L_dom_*), weighted equally (*λ* = 1.0). During backpropagation, the gradient reversal factor “*α*” was dynamically scaled as the training progressed (training progress ratio 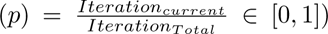 according to the schedule in eq (4):

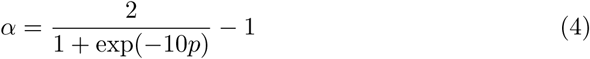

### 2.7 Linear Probing for Domain Separability

Before proceeding to the generalization experiments, we evaluated the extent to which the extracted features retained domain-specific information. To test the domain separability across our representations, we calculated the macro F1-score as a “Dataset Discriminability Score (DDS)” [20]. All samples were pooled, and a Logistic Regression probe was trained using 5-fold stratified cross-validation to perform a multi-class dataset classification (*n* = 7). Nine such classifications were done corresponding to 3 feature types and 3 preprocessings. A binary classification was done as well in the identical experimental conditions to classify the disease labels from the pooled data as a control.

### 2.8 Classifier Training and Evaluation

#### 2.8.1 All vs All Dataset Evaluation

To establish a baseline for generalization, models were evaluated using an All-vs-All cross-dataset strategy prior to the ensemble experiments. In this configuration, a model was trained exclusively on a single ‘source dataset’ and subsequently evaluated on the completely unseen populations of the remaining six datasets. This process was done iteratively until every dataset acted as a source, creating a full evaluation matrix. The trained models from the cross validation evaluated the target datasets to generate independent diagnostic probabilities.

##### Baseline - MLP All vs All Evaluation

The MLP network was trained using the default hyperparameters and regularization constraints defined in the section 2.6.1.

##### DAkk - All vs All Evaluation

The DANN baseline training was done on one dataset at a time, this control experiment isolated the effect of the multiple instance learning without the benefit of multi-source adversarial domain adaptation. To maintain architectural equivalence with subsequent LODO experiments, the DD was retained in this setup. Since, adversarial gradient requires more than one source distributions, evaluating a single domain label yielded a trivial domain loss (*L_dom_*), rendering the adversarial penalty inert. Dynamically calculated balanced class weights were applied to both the clinical and domain loss components to account for heterogeneous label distributions within the source dataset.

#### 2.8.2 Multi-Source Leave-One-Domain-Out (LODO) Ensemble Evaluation

To rigorously test out-of-distribution generalization, models were evaluated using a multi-source leave-one-dataset-out ensemble strategy. In this configuration, six datasets were aggregated to form a diverse training distribution, while the entire population of the seventh dataset was strictly isolated for the final evaluation. This process was repeated iteratively until every dataset functioned as the held-out target. Similar to the all-vs-all setup, upon completing the 5-folds training, the ensemble predicted on the completely unseen held-out dataset.

##### Baseline - MLP LODO Evaluation

The simple MLP served as the non-adversarial ensemble baseline. The pooled source data underwent a stratified 5-fold cross-validation. The training followed the identical structural configurations and training constraints outlined in the section 2.6.1.

##### DAkk - LODO Evaluation

The DANN followed the same structural multi-source pooling strategy to isolate the seventh dataset. Since there were multiple datasets this time in the training pool, it was in this setup that the full architecture of the DANN was working to actively learn the breast cancer specific signals while trying to forget the dataset specific ones.

For a single patient, the TC predicted the binary pathology. Simultaneously, for that exact same patient, the DD executed a 6-way classification to guess the origin dataset of the patient. During backpropagation for that iteration, the TC sent a normal gradient to the FE (teaching it cancer pathology). The DD sent its gradient through the GRL, which was multiplied by −*α*. The FE was thus forced to update its weights to forget whatever dataset-specific noise the discriminator just used to guess the patient’s parent dataset.

#### 2.8.3 Evaluation Metrics and Statistical Analysis

Model performance was primarily quantified using the Area Under the Receiver Operating Characteristic Curve (AUC-ROC) complemented by F1-score, and the Matthews Correlation Coefficient (MCC).

To estimate the uncertainty of the predictions, 95% Confidence Intervals (CIs) were generated using bootstrapping (1000 iterations). The 95% CIs were subsequently derived from the 2.5*^th^* and 97.5*^th^* percentiles of the empirical bootstrap distribution. To assess the statistical significance of the generalization of DANN across batches compared to the generalization of the baseline MLP, two comparisons were made, each using a different sample size and test. For the pooled comparison, scores were aggregated across all seven held-out datasets and three feature extractors within each preprocessing condition, giving *n* = 21 per preprocessing stratum. Normality of the differences was confirmed via the Shapiro-Wilk test at this sample size, and a one-tailed paired t-test was used to test the alternative hypothesis that DANN yields higher AUC-ROC than MLP. For the per-stratum comparison (*n* = 7, one observation per held-out dataset), the Wilcoxon signed-rank test was used instead, as the sample size was too small for the Shapiro-Wilk test to detect departures from normality with adequate power.

To formally test whether the effect of DANN on diagnostic AUC depended on the choice of feature representation that is to test the “interaction effect”, per-stratum AUC gains were defined as ΔAUC = AUC_DANN_ − AUC_MLP_ for each of the 21 LODO strata per feature type (7 held-out datasets × 3 preprocessing conditions). Two complementary analyses were applied to these paired differences. First, three one-sample

Wilcoxon signed-rank tests were used to test whether the median ΔAUC differed from zero within each feature type (H_0_: median ΔAUC = 0), with Bonferroni correction applied across the three simultaneous comparisons. Second, a linear mixed-effects model (LME) was fitted by restricted maximum likelihood (REML) to the full set of 126 AUC observations:

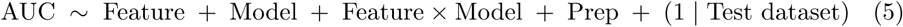

or

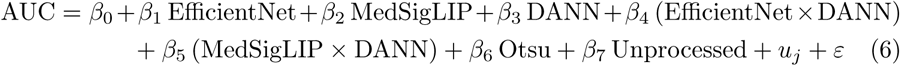

where Feature (Radiomics, EfficientNet, MedSigLIP), Model (MLP, DANN), and Prep (Unprocessed, Otsu, CLAHE+Otsu) were fixed effects, and held-out test dataset 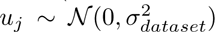 was included as a random intercept for held out dataset *j* to account for baseline AUC differences attributable to dataset-level factors such as scanner type, cohort size, and image acquisition protocol. The 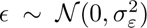 was the residual error. Reference categories were Radiomics (feature), MLP (model), and CLAHE+Otsu (preprocessing). The interaction term “Feature × Model” tests whether DANN’s AUC gain is uniform across representations or differs by feature type. The model was fitted in two independent software environments to assess sensitivity to the choice of inference procedure **(i)** statsmodels (Python 3), which uses asymptotic *z*-tests, and **(ii)** lme4 with lmerTest (R), which applies Satterthwaite-approximated denominator degrees of freedom and *t*-tests. Because only seven held-out datasets were available as random-effect groups, an LME fitting failure under either solver was considered plausible; in that event, the Wilcoxon signed-rank results were designated the primary inference. A visualization of the entire workflow has been presented in the Figure 2

**Fig. 2:**
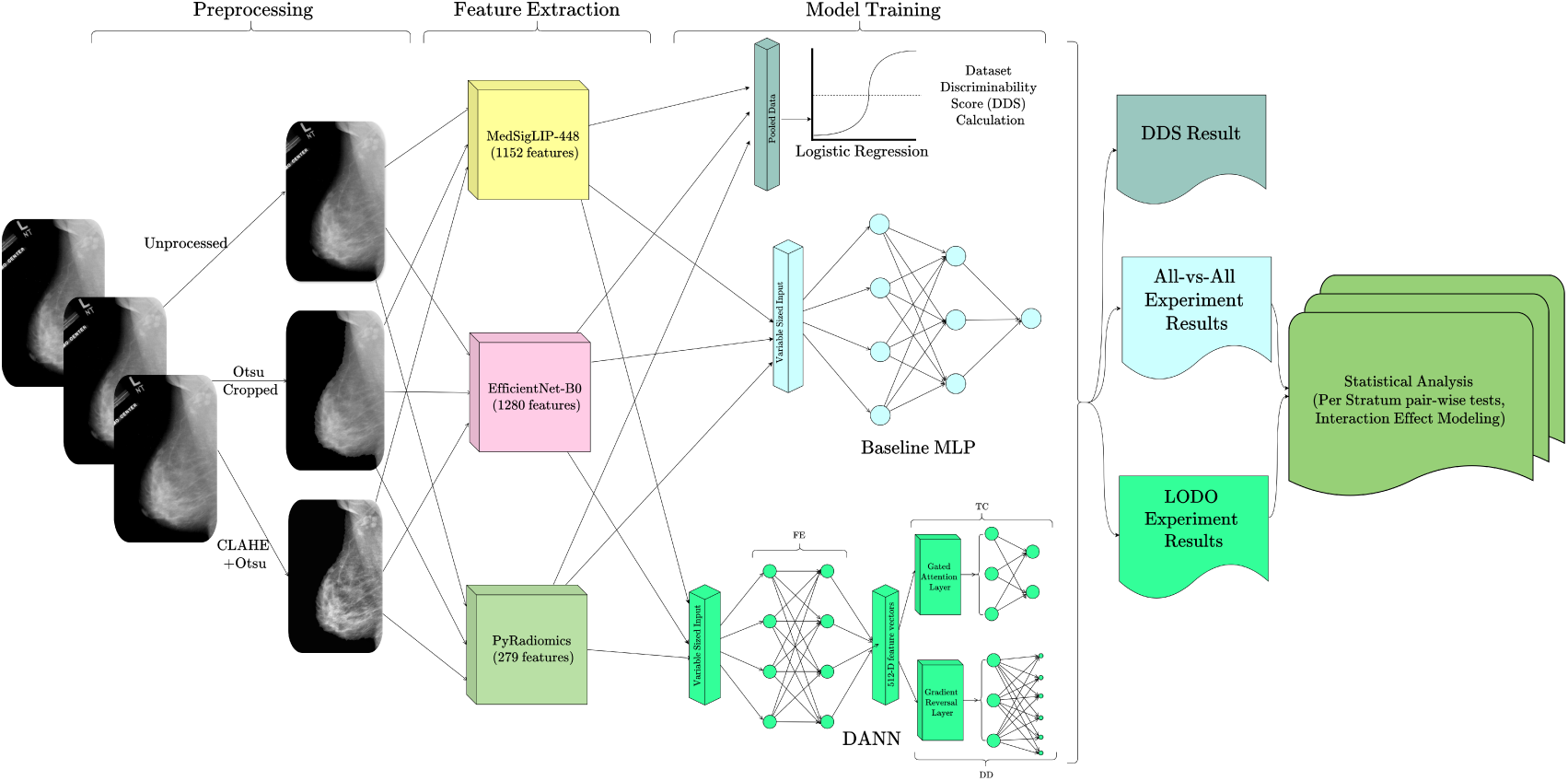
A schematic diagram of the methodology followed in the work

## 3 Results

### 3.1 Preliminary Data Analysis

Detailed characteristics for each dataset, such as the publication year, demographic origin, acquisition modality, machine vendor, and source, are compiled in the Table 1. The precise distribution of patients and image files, categorized by pathology, is presented in Table 2. Figure 3 shows the class distribution and a visualization of geographical origin of the datasets.

**Table 1:** Description of the datasets with their year of publication, demographic origin, acquisition modality, mammography machine vendor and source of download details.

| S.No. | Dataset | Year | Country | Modality | Scanner Vendor | Download source |
| --- | --- | --- | --- | --- | --- | --- |
| 1 | IMDB | 2024 | India | FFDM | Hologic | Indian Biological Images Archive (IBIA) |
| 2 | VinDR-Mammo | 2023 | Vietnam | FFDM | Siemens, IMS, and Planmed | PhysioNet |
| 3 | RSNA | 2022 | US+Australia | FFDM | Hologic, General Electric and Fujifilm | AWS Open Data program |
| 4 | CMMD | 2021 | China | FFDM | General Electric and Siemens | The Cancer Imaging Archive (TCIA) |
| 5 | CBIS-DDSM | 2016 | United States | SF | Details unavailable | The Cancer Imaging Archive (TCIA) |
| 6 | INbreast | 2012 | Portugal | FFDM | Siemens | Kaggle |
| 7 | MIAS | 1994 | United Kingdom | SF | Details unavailable | Apollo - University of Cambridge Repository |

**Table 2:** Number of patients and image files in each of the datasets utilized on the basis of their cancer pathology.

| S.No. | Dataset | Image files |  |  | Patients |  |  |
| --- | --- | --- | --- | --- | --- | --- | --- |
|  |  | Benign | Malignant | Total | Benign | Malignant | Total |
| 1 | IMDB | 3277 | 300 | 3577 | 510 | 73 | 583 |
| 2 | VinDR-Mammo | 2860 | 988 | 3848 | 481 | 481 | 962 |
| 3 | RSNA | 5014 | 1069 | 6083 | 1312 | 443 | 1755 |
| 4 | CMMD | 1084 | 1130 | 2214 | 473 | 553 | 1026 |
| 5 | CBIS-DDSM | 1728 | 1375 | 3103 | 814 | 752 | 1566 |
| 6 | INbreast | 310 | 100 | 410 | 59 | 49 | 108 |
| 7 | MIAS | 269 | 53 | 322 | 110 | 51 | 161 |
| 8 | Total | 14542 | 5015 | 19557 | 3759 | 2402 | 6161 |

**Fig. 3:**
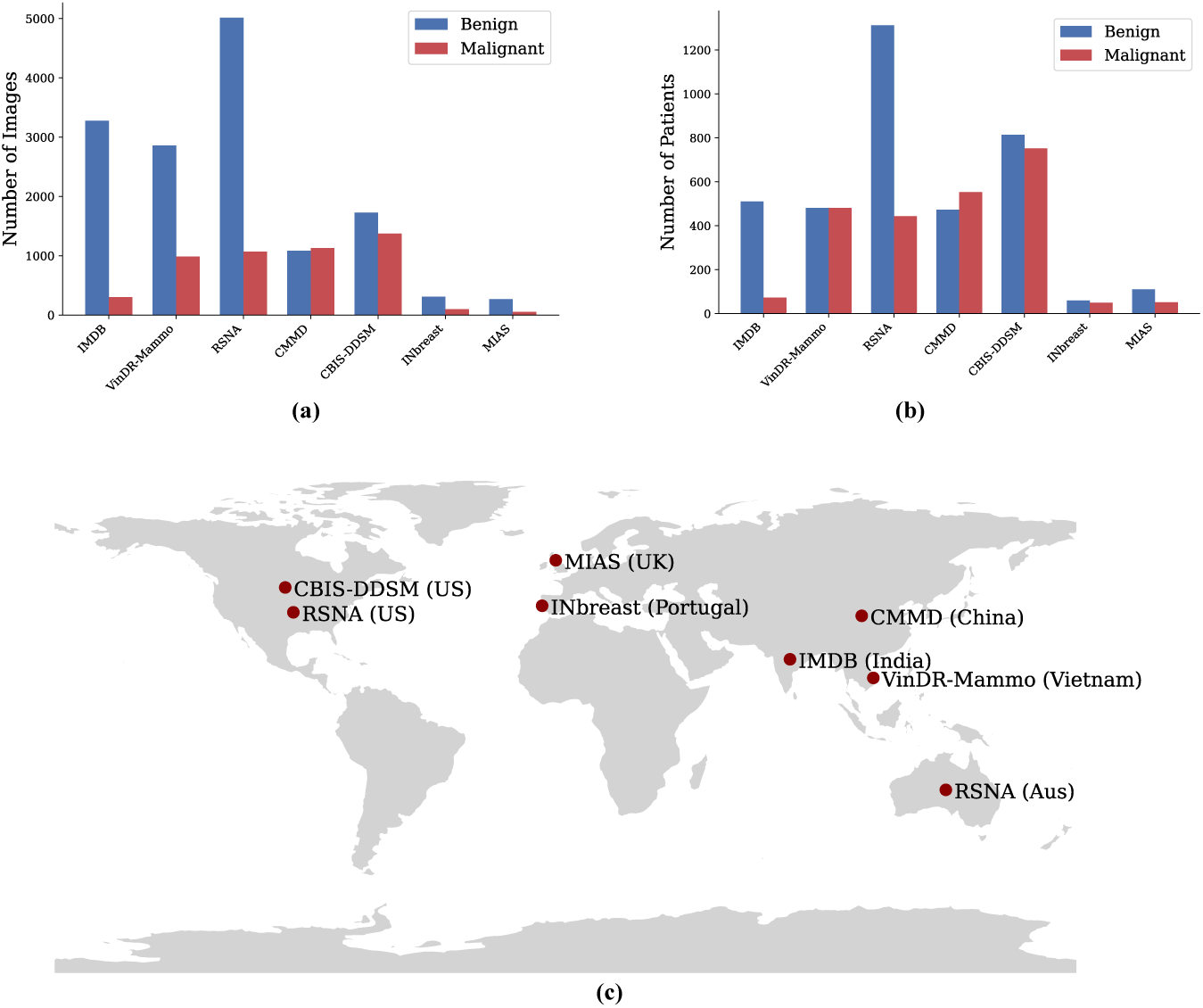
Bar plot (a) shows the class distribution (Benign vs Malignant) in the datasets in terms of number of images while that in terms of number of patients is shown in the plot (b). Figure (c) shows a schematic visualization of the geographical origin of the datasets.

### 3.2 Domain Separability of Representations

In the dataset discriminability experiment, the DDS was calculated revealing that in EfficientNet and MedSigLIP features the clinical and dataset specific signals were deeply entangled. Radiomics however, had no such domain specific patterns in the representations it generated. In the Figure 4a we can see in terms of macro f1 scores, how highly separable the data cohort is with EfficientNet and MedSigLIP features, each yielding a mean macro f1 score of 0.9687±0.005 and 0.9622±0.001 respectively across preprocessings. Radiomics on the other hand was able to successfully erase the dataset specific signatures and the mean macro f1 stayed around 0.1166 ± 0.01).

**Fig. 4:**
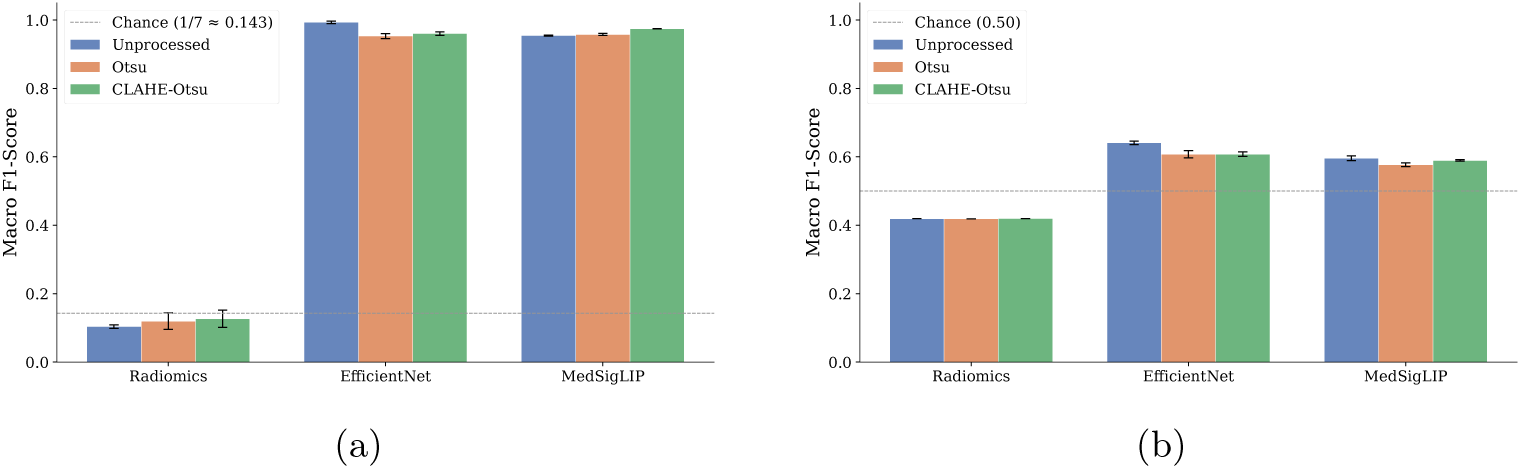
(a) Macro f1 as the Dataset Discriminability Score (DDS) [20] evaluating the linear separability of domain-specific information across feature representations and preprocessing conditions. (b) Macro f1 for the label classification using the same linear probe in identical experimental conditions.

As a control experiment, we tried predicting the disease labels from the same pooled cohort and the linear probe, the mean macro f1 for EfficientNet and MedSigLIP were 0.6187 ± 0.007 and 0.5873 ± 0.004 respectively and the same for radiomic features was 0.4190±0.48×10^−5^. Although, the EfficientNet and MedSigLIP features provided some gain in performance, radiomics stayed much below the random chance and as compared to the linear separability of datasets, that of disease labels in identical conditions completely failed. Figure 4b shows the results.

### 3.3 Baseline MLP Performance

The 3-layer MLP baseline, evaluated under the All-vs-All framework, achieved an overall aggregate AUC-ROC of 0.539 [95% CI: 0.53, 0.54]. The mean in-domain AUC-ROC was 0.630 [95% CI: 0.608, 0.651]. Cross-domain performance dropped to 0.524 [95% CI: 0.521, 0.528], a collapse toward random chance that held across all preprocessing and feature configurations. Interestingly, unprocessed images consistently outperformed both Otsu-cropped and CLAHE+Otsu-processed inputs across feature types, the opposite of what preprocessing pipelines are conventionally expected to achieve. However, even the **best-performing configuration, “unprocessed images paired with MedSigLIP features”**, could not bridge the gap between in-domain and cross-domain performance; it produced an in-domain mean AUC-ROC of 0.728 [95% CI: 0.666, 0.791] against a cross-domain mean of 0.573 [95% CI: 0.563, 0.584].

The performance violin plots in Figure 5a reflect the in-domain vs cross-domain gap across all three preprocessing conditions and all three feature extractors. The transferability matrix in Figure 5c also validates this pattern.

**Fig. 5:**
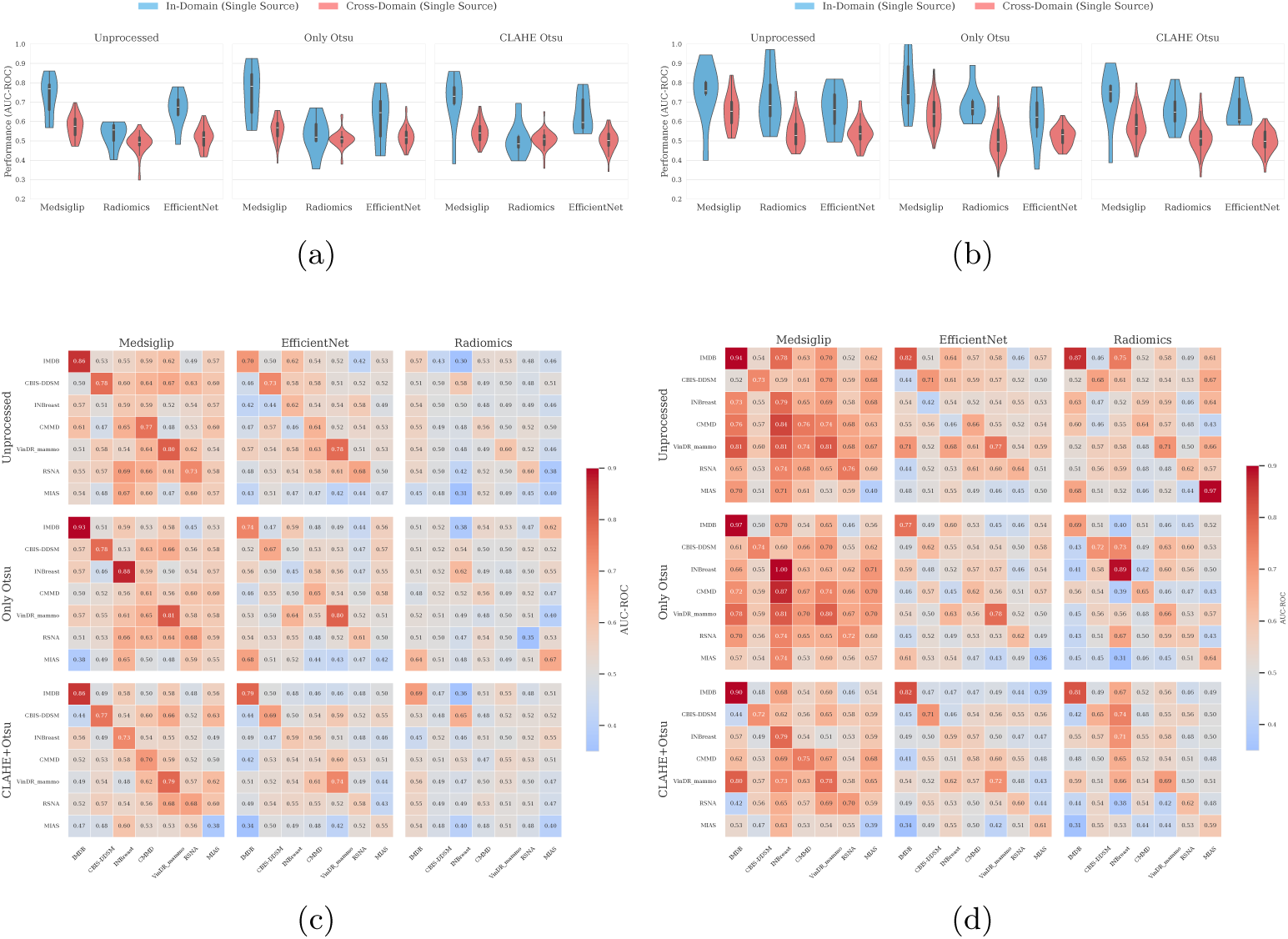
Combined performance metrics for MLP and DANN architectures. (a) Violin plots showing in-domain versus cross-domain generalization performance for the 3-layer MLP baseline, stratified by preprocessing and feature extractor. (b) Violin plots showing the in-domain versus cross-domain generalization performance for the DANN architecture (discriminator disabled), stratified by preprocessing and feature extractor. (c) Heatmaps showing MLP baseline performance matrices. Diagonal cells represent in-domain performance; off-diagonal cells represent cross-domain transfer and fall below diagonal values across every source-target pair without exception. (d) Heatmap showing the performance matrix in the All-vs-All DANN experiment (domain discriminator disabled). Diagonal cells represent in-domain test splits; off-diagonal cells represent cross-domain transfer.

### 3.4 MLP LODO Ensemble Approach and Partial Recovery

Replacing the all vs all baseline training scheme with a Leave-One-Dataset-Out (LODO) ensemble partially recovered cross-domain performance. The MLP LODO ensemble reached an aggregate mean AUC-ROC of 0.545 [95% CI: 0.537, 0.554], compared to 0.524 [95% CI: 0.521, 0.528] for the cross-domain All-vs-All MLP baseline (LODO ensemble results are inherently cross-domain, so instead of comparing them with aggregate, they have been compared with cross-domain results), a modest but consistent gain across configurations. Under the reference configuration of unprocessed images with MedSigLIP features, the ensemble reached a mean AUC-ROC of 0.598 [95% CI: 0.574, 0.622]. MedSigLIP and EfficientNet features both generalized better than Radiomics under the ensemble scheme, with Radiomics performance remaining near the chance line regardless of preprocessing as reflected in the Figure 6a.

**Fig. 6:**
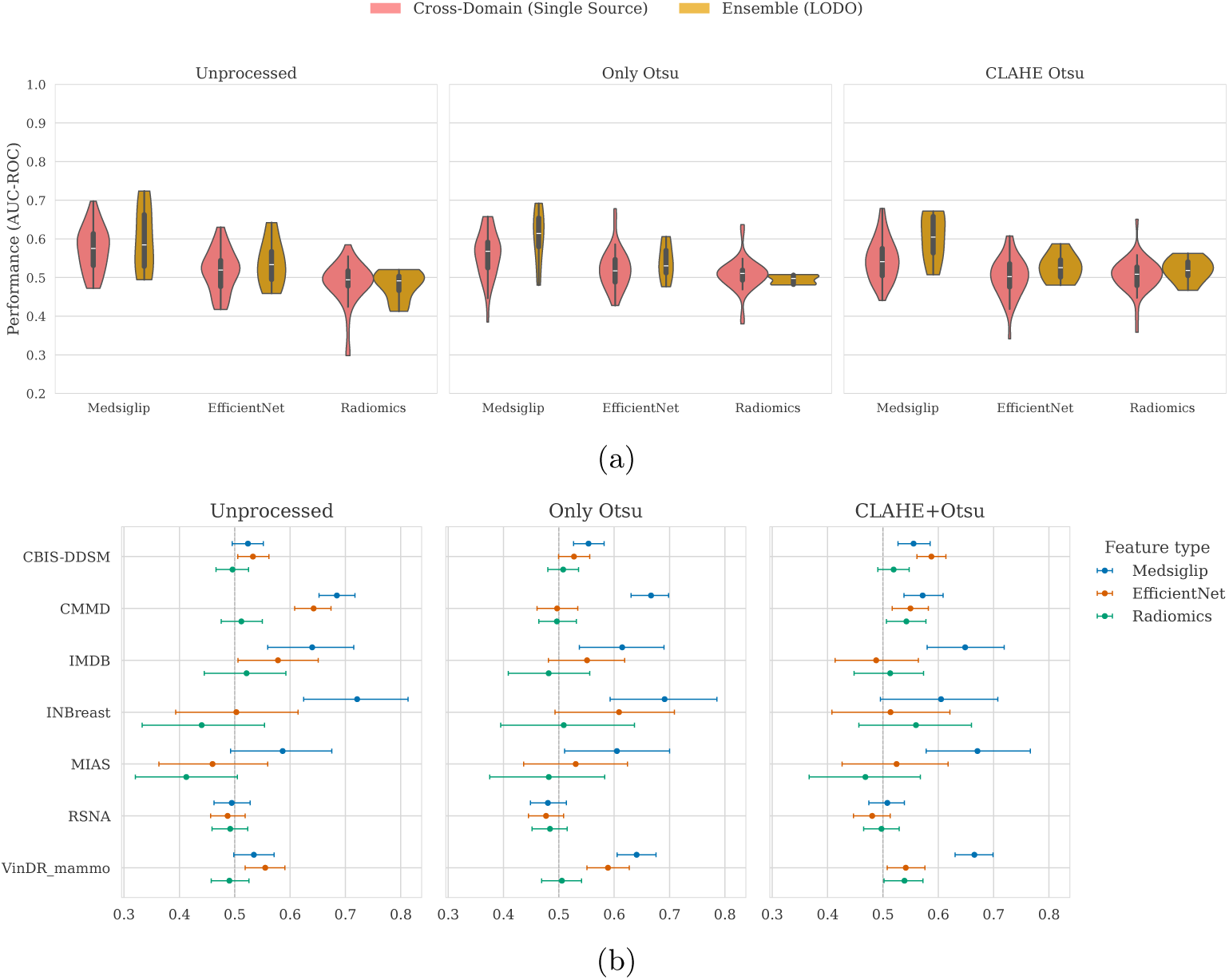
(a) The cross-domain versus ensemble generalization performance for the 3-layer MLP, stratified by preprocessing and features. The Forest plots in (b) AUC-ROC performance (with 95% C.I.) on the x axis by held-out datasets on the y axis for the MLP LODO ensemble under the three preprocessing conditions.

Per-dataset forest plots in Figure 6b show that this aggregate gain was not uniform. Held-out performance varied substantially by dataset, with some targets benefiting clearly from ensemble training while others remained at or below the random chance. Under the reference configuration, per-dataset AUC-ROC ranged from 0.724 [95% CI: 0.625, 0.813] for INBreast to 0.494 [95% CI: 0.462, 0.528] for RSNA.

The ensemble too did not close the performance gap, and reached a limit at approximately AUC-ROC ≈ 0.7, even for the best configurations of feature, preprocessing and dataset. The limit in performance has a straightforward cause; LODO averaging pools source datasets but does not constrain what the model learns during training. Each constituent model still optimizes for classification without any pressure to ignore dataset-specific variation, so the ensemble reproduces whatever batch signal each individual model has absorbed. Suppressing that signal during training, rather than averaging over it after, is what the domain-adversarial approach is designed to do, which motivated the same.

### 3.5 Generalizability of DANN (Without GRL) vs That of MLP

With the Domain Discriminator disabled, the DANN architecture evaluated under the same All-vs-All conditions as previous experiments, already generalized better than the MLP baseline. Over all aggregate DANN All-vs-All AUC-ROC was 0.575 [95% CI: 0.571, 0.579] compared to baseline (MLP) All-vs-All aggregate of 0.539 [95% CI: 0.535, 0.543]. In-domain performance was 0.693 [95% CI: 0.674, 0.713], cross-domain AUC-ROC was 0.556 [95% CI: 0.552, 0.559], compared to 0.630 [95% CI: 0.608, 0.651] and 0.524 [95% CI: 0.521, 0.528] for the MLP. Under the reference configuration of unprocessed images with MedSigLIP features, in-domain AUC-ROC was 0.742 [95% CI: 0.683, 0.801] and cross-domain AUC-ROC was 0.652 [95% CI: 0.643, 0.662], with respect to performances in MLP LODO (0.59) and MLP All-vs-All cross-domain (0.57). The preprocessing ordering observed in the MLP experiments “Unprocessed > Only-Otsu > CLAHE+Otsu” held here as well, as did the feature type ordering, with MedSigLIP leading EfficientNet and Radiomics across all conditions. Figure 5b shows the performance violin plots per stratum and Figure 5d shows the transferability matrix across in domain and cross domain in the all vs all experiment.

### 3.6 DANN LODO Ensemble: Optimal Configuration and Significance

Applying the full DANN architecture with the Gradient Reversal Layer (GRL) and Domain Discriminator (DD) active, under LODO cross-validation produced the best generalization results of all evaluated configurations. The aggregate mean AUC-ROC was 0.575 [95% CI: 0.567, 0.583] compared to DANN All-vs-All cross domain performance (0.55), MLP LODO performance (0.54) and MLP All-vs-All cross-domain performance (0.52). Under the reference configuration of unprocessed images with MedSigLIP features, the mean AUC-ROC reached 0.685 [95% CI: 0.663, 0.706] against DANN All-vs-All cross domain=0.652 [95% CI: 0.643, 0.662], MLP LODO=0.598 [95% CI: 0.574, 0.622] and MLP All-vs-All=0.573 [95% CI: 0.563, 0.584]. Per-dataset performance under this configuration ranged from 0.507 [95% CI: 0.475, 0.540] for RSNA (most difficult to generalize target across all experiments) to 0.871 [95% CI: 0.806, 0.929] for INBreast shown in Figure 7a.

**Fig. 7:**
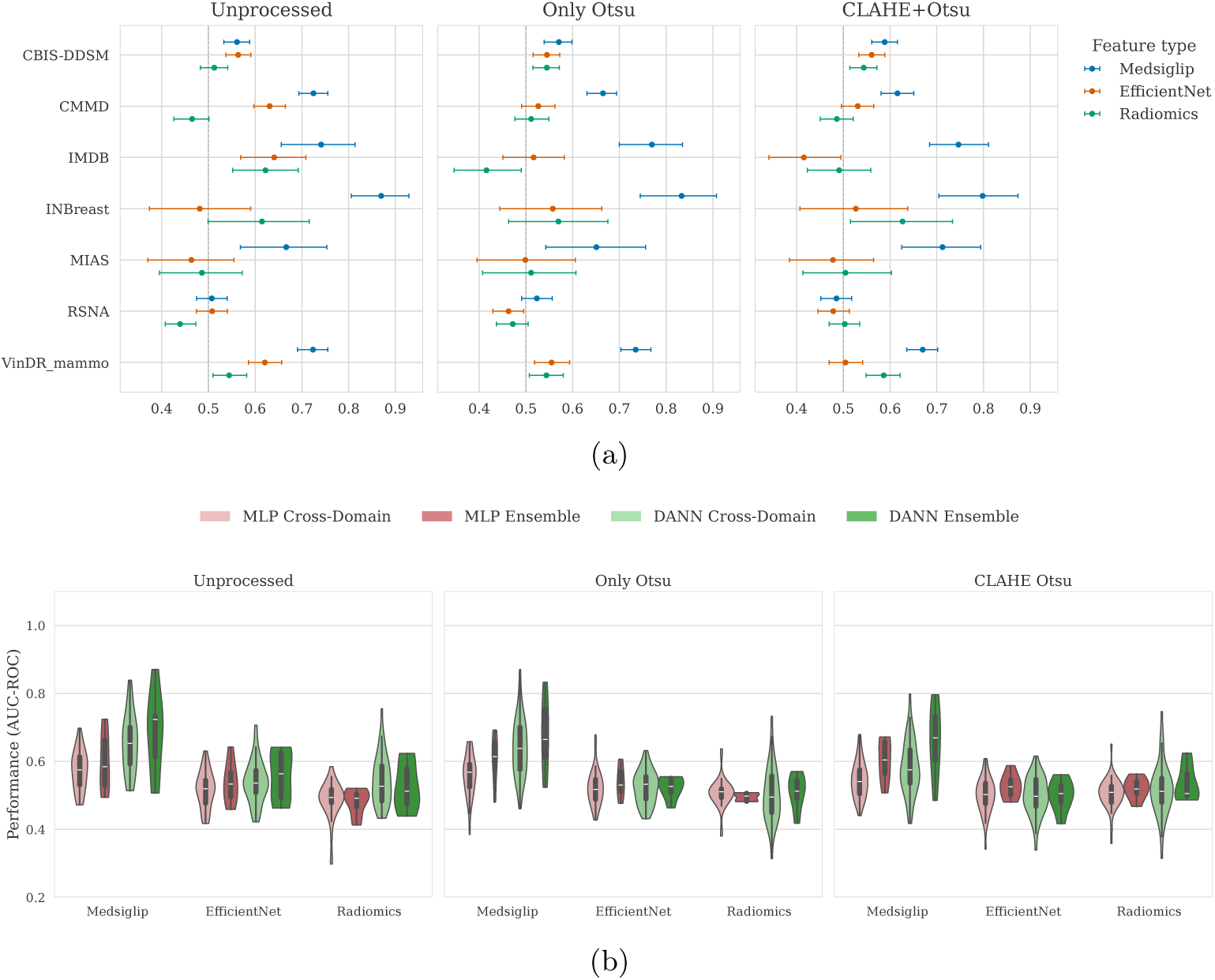
(a) AUC-ROC by held-out dataset for the DANN LODO ensemble (with 95% C.I.). Each panel corresponds to one preprocessing condition; feature types are shown as dodged points in 3 colors within each dataset. (b) Plot shows cross-domain versus ensemble generalization across all four evaluation regimes (MLP all vs all cross-domain, MLP LODO ensemble, DANN all vs all cross-domain, DANN LODO ensemble), stratified by preprocessing and features.

The Figure 7b places these results in the context of all four evaluation regimes MLP cross-domain, MLP LODO ensemble, DANN cross-domain, and DANN LODO ensemble; separated by preprocessing conditions and feature types. The progression from MLP all vs all cross-domain (light pink) to DANN LODO ensemble (dark green) is most pronounced for MedSigLIP features from unprocessed images, where the median shifts upward and the distribution narrows considerably. For EfficientNet and Radiomics the four regimes converge near chance, confirming that the gains are specific to MedSigLIP and do not generalize to the other two feature types under the current architecture.

### 3.7 Statistical Test Results

#### 3.7.1 Stratified Pair-wise Analysis

The pooled paired t-test (n=21) shown in Table 3 confirmed that DANN LODO ensemble performance exceeded MLP LODO ensemble performance for MedSigLIP features; p value with Bonferroni correction (*p_Bonf._*= 7.34 × 10^−5^ against an adjusted *α*=0.0167). For Radiomics, the raw p value reached nominal significance (p = 2.92 × 10^−2^, p < 0.05), but as the correction was applied this effect vanished (*p_Bonf._*=8.74 × 10^−2^). DANN did not significantly outperform MLP for EfficientNet features in the pooled comparison (*p_Bonf._*=1); the negative t-statistics for Efficient-Net indicates DANN slightly underperformed MLP in those conditions, though not significantly.

**Table 3:** Paired One-tailed t-test results (Bonferroni corrected) on the alternate hypothesis (H1) “DANN LODO Ensemble provides better cross domain generalization than MLP LODO Ensemble in terms of bootstrapped mean AUC-ROC” and null hypothesis (H0) “no betterment in generalization due to DANN LODO Ensemble as compared to MLP LODO Ensemble”. Done for AUC-ROC values aggregated across the 3 preprocessing conditions.

| Feature Type | n | Mean MLP | Mean DANN | t statistic | $p_{Bonf.}$ | Adjusted $\alpha$ | $\beta$ | Significant? |
| --- | --- | --- | --- | --- | --- | --- | --- | --- |
| **MedSigLIP** | **21** | **0.603** | **0.674** | **5.1481** | **$7.3422 \times 10^{-5}$** | **0.0167** | **0.00332** | **Yes** |
| Radiomics | 21 | 0.499 | 0.524 | 2.0084 | $8.7452 \times 10^{-2}$ | 0.0167 | 0.59217 | No |
| EfficientNet | 21 | 0.534 | 0.527 | -0.9433 | 1 | 0.0167 | 0.99874 | No |

Done for each preprocessing condition and feature type, Wilcoxon signed-rank test (n=7, one observation per held-out dataset) results are reported in Table 4 and Figure 8. For MedSigLIP, DANN significantly exceeded MLP in all three preprocessing strata: Unprocessed (p = 7.81×10^−3^, p < 0.01), Only-Otsu (p = 1.56×10^−2^, p < 0.05), and CLAHE+Otsu (p = 2.34 × 10^−2^, p < 0.05). Neither Radiomics nor EfficientNet reached significance in any preprocessing stratum, consistent with the pooled results.

**Table 4:** Wilcoxon Signed-Rank Test Results on the alternate hypothesis (H1) “DANN LODO Ensemble provides better cross domain generalization than MLP LODO Ensemble in terms of bootstrapped mean AUC-ROC” and null hypothesis (H0) “no betterment in generalization due to DANN LODO Ensemble as compared to MLP LODO Ensemble”. Done for AUC-ROC values per preprocessing condition and feature type.

| Feature Type | Preprocessing | n | Mean MLP | Mean DANN | W statistic | p-value | Rank-biserial r | Significant? |
| --- | --- | --- | --- | --- | --- | --- | --- | --- |
| MedSigLIP | Unprocessed | 7 | 0.598 | 0.685 | 28.00 | $7.8125 \times 10^{-3}$ | 0.0000 | Yes** |
| MedSigLIP | Only Otsu | 7 | 0.607 | 0.678 | 27.00 | $1.5625 \times 10^{-2}$ | 0.0357 | Yes* |
| MedSigLIP | CLAHE+Otsu | 7 | 0.603 | 0.659 | 26.00 | $2.3428 \times 10^{-2}$ | 0.0714 | Yes* |
| Radiomics | Unprocessed | 7 | 0.481 | 0.526 | 23.00 | $7.8125 \times 10^{-2}$ | 0.1786 | No |
| Radiomics | Only Otsu | 7 | 0.495 | 0.510 | 20.00 | $1.8750 \times 10^{-1}$ | 0.2857 | No |
| Radiomics | CLAHE+Otsu | 7 | 0.520 | 0.535 | 20.00 | $1.8750 \times 10^{-1}$ | 0.2857 | No |
| EfficientNet | Unprocessed | 7 | 0.537 | 0.558 | 22.00 | $1.0938 \times 10^{-1}$ | 0.2143 | No |
| EfficientNet | Only Otsu | 7 | 0.540 | 0.523 | 05.00 | $9.4531 \times 10^{-1}$ | 0.8214 | No |
| EfficientNet | CLAHE+Otsu | 7 | 0.526 | 0.499 | 02.00 | $9.8438 \times 10^{-1}$ | 0.9286 | No |

Significance codes: \*\* $p < 0.01$ ; \* $p < 0.05$ .

**Fig. 8:**
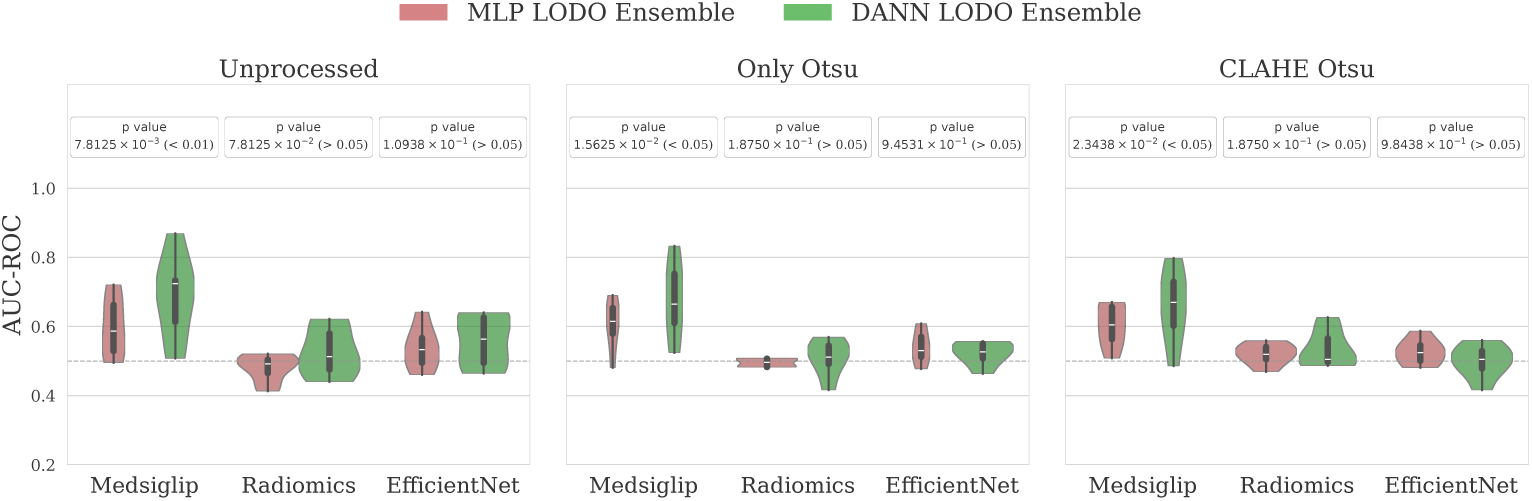
MLP versus DANN LODO ensemble AUC-ROC distributions stratified by preprocessing condition and feature type, with per-stratum p-values annotated.

A Bonferroni correction was omitted for this per-stratum analysis due to the severe power starvation inherent to non-parametric tests at *n* = 7;. Moreover, the primary evidence of significance of DANN LODO strategy being better than the MLP LODO was already there from the t-test, and this stratified analysis was done to detect the edge cases and localized behavior.

#### 3.7.2 Global Interaction Analysis

The linear mixed effect (LME) model converged under both implementations with identical coefficient estimates; the random intercept variance was 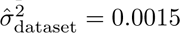 and the residual variance was 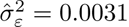. The interaction term for MedSigLIP × DANN was 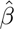_5_ = +0.046 (SE = 0.024). This resulted in *z* = 1.90, *p* = 0.058 under the asymptotic *z*-test and *t*(112) = 1.90, *p* = 0.060 using Satterthwaite’s degrees of freedom approximation. The interaction term for EfficientNet × DANN was 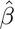_4_ = −0.033 (SE = 0.024; *p* = 0.185). Preprocessing condition had no significant effect on AUC under either implementation (all *p >* 0.55).

The Wilcoxon signed-rank test done for the interaction effect showed that DANN produced a significant median AUC gain only for MedSigLIP (median ΔAUC = +0.044, *p*_Bonferroni_ = 0.00004 (*<* 0.001)). For Radiomics (median ΔAUC = +0.029; *p*_Bonferroni_ = 0.210) and EfficientNet (median ΔAUC = −0.014; *p*_Bonferroni_ = 0.772), no significant gain was detected. Full results are reported in Table 5.

**Table 5:** Representation × DANN interaction analysis. *Panel A*: per-feature one-sample Wilcoxon signed-rank tests on ΔAUC (= AUC_DANN_ − AUC_MLP_), Bonferroni-corrected (*n* = 21 per feature type). *Panel B* : LME fixed-effect coefficients, asymptotic *z*-tests (statsmodels, Python; REML). *Panel C* : LME fixed-effect coefficients, Satterthwaite *t*-tests (lme4/lmerTest, R; REML). Reference categories: Radiomics (feature), MLP (model), CLAHE+Otsu (preprocessing).

| *(A) One-sample Wilcoxon signed-rank test ($H_0$: median $\Delta\text{AUC} = 0$)* |  |  |  |  |  |
| --- | --- | --- | --- | --- | --- |
| Feature type | Median $\Delta\text{AUC}$ | Mean $\Delta\text{AUC}$ | $W$ | $p_{\text{Bonf.}}$ | Sig. |
| Radiomics | +0.029 | +0.025 | 63.0 | 0.210 | No |
| EfficientNet | -0.014 | -0.007 | 82.0 | 0.772 | No |
| **MedSigLIP** | **+0.044** | **+0.071** | **6.0** | **0.00004** | **Yes\*\*\*** |
| *(B) LME fixed effects – asymptotic $z$-tests (statsmodels, Python; REML)* |  |  |  |  |  |
| Term | $\hat{\beta}$ | SE | $z$ | $p$ | Sig. |
| Intercept | 0.496 | 0.020 | 24.578 | $< 0.001$ | Yes*** |
| Feature: EfficientNet | +0.036 | 0.017 | 2.086 | 0.037 | Yes* |
| **Feature: MedSigLIP** | **+0.104** | **0.017** | **6.061** | $< 0.001$ | **Yes\*\*\*** |
| Model: DANN | +0.025 | 0.017 | 1.455 | 0.146 | No |
| Prep: Otsu | +0.002 | 0.012 | 0.149 | 0.882 | No |
| Prep: Unprocessed | +0.007 | 0.012 | 0.576 | 0.564 | No |
| EfficientNet × DANN | -0.033 | 0.024 | -1.335 | 0.182 | No |
| **MedSigLIP × DANN** | **+0.046** | **0.024** | **1.897** | **0.058** | . |
| *(C) LME fixed effects – Satterthwaite $t$-tests (lme4/lmerTest, R; REML)* |  |  |  |  |  |
| Term | $\hat{\beta}$ | SE | $t$ | $p$ | Sig. |
| Intercept (df = 17.7) | 0.503 | 0.020 | 24.927 | $< 0.001$ | Yes*** |
| Feature: EfficientNet (df = 112) | +0.036 | 0.017 | 2.086 | 0.039 | Yes* |
| **Feature: MedSigLIP (df = 112)** | **+0.104** | **0.017** | **6.061** | **$1.87 \times 10^{-8}$** | **Yes\*\*\*** |
| Model: DANN (df = 112) | +0.025 | 0.017 | 1.455 | 0.149 | No |
| Prep: Otsu (df = 112) | -0.005 | 0.012 | -0.428 | 0.670 | No |
| Prep: CLAHE+Otsu (df = 112) | -0.007 | 0.012 | -0.576 | 0.566 | No |
| EfficientNet × DANN (df = 112) | -0.033 | 0.024 | -1.335 | 0.185 | No |
| **MedSigLIP × DANN (df = 112)** | **+0.046** | **0.024** | **1.897** | **0.060** | . |

Significance codes: \*\*\* $p < 0.001$ ; \*\* $p < 0.01$ ; \* $p < 0.05$ ; . $p \geq 0.05$ ; No $p \geq 0.10$ . LME random-effect variance: held-out dataset $\sigma_{\text{dataset}}^2 = 0.0015$ , residual $\sigma_{\varepsilon}^2 = 0.0031$ . Wilcoxon $p$ -values are Bonferroni-corrected across three simultaneous tests ( $\alpha = 0.0167$ ). Satterthwaite degrees of freedom for the intercept reflect estimation primarily from between-group variation ( $k = 7$ groups).

While the LME interaction term is marginally non-significant, the Wilcoxon estimation indicates strict directional superiority (*p* = 4 × 10^−5^). This discrepancy occurs because the LME model applies continuous parametric assumptions and partitions variance to the random intercepts, whereas rank based Wilcoxon test evaluates consistent fold wise improvements independently of magnitude or distribution tails.

Taken together, the results point to one configuration as the recommended approach for cross-population mammography classification: **DANN LODO ensemble training using MedSigLIP features**. MedSigLIP-448 features consistently achieved above-chance AUC-ROC across all preprocessing conditions (DANN mean: 0.659-0.685), while Radiomics and EfficientNet-B0 remained near chance (means<0.56 in all conditions), suggesting that large-scale medical vision pretrained models are better suited for cross-domain mammography generalization as they contain one of the highest amount of dataset specific signatures as evident from the DDS and statistical tests together with the interaction effect modeling to know the interaction effect between different preprocessings, feature types and classifier models. This is precisely why the DANN failed to generalize the radiomic features because they were statistical in nature and never had any dataset bias.

## 4 Discussion and Conclusion

In this study seven mammogram datasets were used to examine the three successive standard preprocessing stages, three different feature extraction strategies, two classifier models and two different training strategies with the goal of minimizing the batch-effects present in the mammograms and maximizing the model generalizability across datasets of varying origin and nature for breast cancer detection. Apart from the biases introduced during image acquisition and the ones coming directly from differences in population, anatomy etc there are other factors which can contribute towards stopping a model from generalizing. The bias mitigation during acquisition becomes a research question for clinicians, physicists and scientists working in the field of nuclear medicine, and we looked at this problem purely from a computational perspective as such a solution can be much more easily deployed in real world conditions as compared to a pioneering hardware solution.

Thus, we started by systematically testing each of the stages which are there in mammogram and biomedical image analysis in general. The three cohorts of the data on basis of preprocessing (unprocessed, Otsu cropped and CLAHE+Otsu cropped) underwent feature extraction with three different methods prevalent in the field (PyRadiomics; radiomic features, EfficientNet-B0; CNN features and MedSigLIP-448; ViT embeddings) each being a representative of the technological advancement over it’s previous one. Once the features were ready for training, we chose to work with Domain Adversarial Neural Network as our primary classifier due to its promising architecture precisely for the kind of domain-generalization task we sought to do, a simple three layer MLP was chosen as a control model. However, even before starting the classification experiments, we needed to see which of the features carry most amount of dataset signature such that a linear probe is able to identify the dataset from a pool of randomized samples (patient ids), for this we trained a Logistic Regression model for doing a multi-class classification (n=7 for seven datasets) and the macro f1 scores were recorded as “Dataset Discriminability Score (DDS)” in accordance with recent literature. The results were interesting but somewhat expected, as this classification showed that the convolutional and the ViT features were picking up the dataset specific signals and entangling them with clinical signals so strongly that the Logistic Regression was able to classify the datasets with mean macro f1 going above 0.96 but was completely failing to do so when trying to classify the disease labels. Radiomics on the other hand was erasing the dataset level information successfully but was removing the important clinical information as well as the classifier stayed below the random chance line in both the classifications. This suggests that the entanglement of the dataset specific and clinical information is such that if we try to remove one of them, the other gets lost as well leaving us with noise. The CNN and ViT features however showed some positive trend towards the generalizability as even though they had more domain specific signatures but had more pathology information as well.

To move towards the actual domain generalization experiments, we started by training a three layer MLP in an all-vs-all design. Through this experiment we wanted to quantify the “single-source training” generalization baseline. The best generalization was seen with MedSigLIP features coming from unprocessed images. After this we trained the DANN model following the same experimental design and the trend remained consistent as before with MedSigLIP features from unprocessed images giving the best generalization among the other preprocessings and features. Here, as we were doing a “single-source training” and “multi target evaluation” the GRL of DANN was rendered inert as it needs at least two source domains for being active. Thus, the DANN in this setup did a multi-instance learning with gated attention to focus more on the patches containing the most disease specific signal and improved the performance marginally with respect to MLP all-vs-all. As we had hit a limit on performance improvement, the next step was to switch to ensemble trainings. Since we working with a decent number of datasets, we did “Leave One Dataset Out (LODO)” ensemble trainings with both the MLP and the DANN. The MLP LODO results stood between MLP all-vs-all and the DANN all-vs-all results, making it clear that the MedSigLIP features were performing the best in terms of generalization and the DANN without GRL itself was superior to the MLP even in single-source training and evaluation. The final training was done with DANN in LODO configuration. In this setup, the GRL was working actively to dis-entangle the dataset specific and cancer specific signals, and this setup gave us the best results which was a mean cross-domain AUC-ROC of 0.685 [95% CI: 0.663, 0.706] with the best performing, MedSigLIP features coming from unprocessed images. The best performing held out dataset was INBreast (0.871 [95% CI: 0.806, 0.929]) and worst performance was of RSNA dataset which stood around the random chance. This difficult to generalize to behavior of RSNA has been documented in published literature as well.

From all the experimentation a conclusion was derived that the DANN LODO training on MedSigLIP features coming from unprocessed images is the way to go to get the best cross-cohort generalization. To test this formally, pairwise parametric and non-parametric tests were done in which this hypothesis held true with substantial significance. In the study however, since initial stages, multiple variables were involved, thus we decided to test if any interaction is going on among them, and if yes then how is it affecting the eventual generalization performance. A “Linear Mixed Effect (LME)” model was used alongside a non-parametric estimation using the Wilcoxon Signed rank test. The LME modeling itself was done in two different software environments in Python3 (asymptotic z test) and R (Satterthwaite t test) to get the best opinion. In both the implementations it was established that the MedSigLIP provides a far better representation as compared to radiomics and EfficientNet towards generalizability but DANN’s contribution was insignificant as compared to MLP. The effect of MedSigLIP when used with DANN was directionally consistent but missed the significance level marginally in the LME modeling. The real picture emerged when we did the Wilcoxon Signed rank estimation where after doing the Bonferroni correction, the AUC_DANN_ − AUC_MLP_ effect was highly significant (4 × 10^−5^ against an adjusted significance level of 0.0167) for MedSigLIP features whereas it was insignificant for other features. In all the post-hoc tests, the significant effect of unprocessed images or the effect of preprcessings was ruled out in general.

This gave us few clean conclusions **(i)** Domain generalization for mammograms is a problem which can be solved computationally by making careful choices during feature extraction and classifier training. **(ii)** The choice of feature extractor model is crucial as the amount of usable information can vary with varying models. MedSigLIP-448 is one open-source vision-language foundation model which can help us in getting the best cross-cohort generalization performance for mammography out of the box (without any fine tuning) even though mammograms lack from its training data. **(iii)** Choice of the classifier model while dealing with a domain generalization problem is equally important as the feature extractor choice, and architectures like that of DANN solve that problem very elegantly and with superior performance. **(iv)** Surprisingly, the effect of preprocessing of the images is insignificant overall for the task in question. **(v)** Dataset identity traveling in the FM based image representations is not necessarily a thing which we need to be cautious of, as we have seen that when we work with architectures like DANN, it is able to disentangle the dataset and clinical signals very well in the representations which had a very high batch specific signal but had a very rich disease specific signal as well.

## 5 Future Directions

After exploring different methodologies and establishing the superiority of Domain Adversarial Neural Network (DANN) in cross-cohort mammography generalization using MedSigLIP features, as a logical next step, we can move towards fine tuning the MedSigLIP-448 model on diverse mammography datasets, as they are the one radiological modality which are missing from its training data and we have done a “Zero-shot” transfer of learning in this study which looks promising. Also, we can then benchmark the fine-tuned MedSigLIP against different mammography specific foundation models like MammoCLIP [17], mamaDINO [39] etc. This will be interesting to see as in our study we have used whole mammograms devoid of “Region of Interest (ROI)” annotation for tumors, while most of the published studies have relied on the ROI information while training their models.

## Declarations

### • Data availability

All the datasets utilized in the study are open-access and are freely available to download from their dedicated portals. All the details regarding their acquisition have been discussed in the section 2.1 and the associated literature has been duly cited.

### • Code availability

All the scripts developed during this work are available in our GitHub repo github.com/Sciwhylab. Codes will be kept private till publication.

### • Author contribution

SA Conceptualized and coordinated the project, including final manuscript editing and study design. AKP curated the data, developed the methodology and performed experiments.

## Acknowledgments

AKP would like to thank the entire SciWhyLab group, current and former members, for creating a healthy, vibrant and fun environment to be in and his mentor SA for believing in him.

